# YeastNet: Deep Learning Enabled Accurate Segmentation of Budding Yeast Cells in Bright-field Microscopy

**DOI:** 10.1101/2020.11.30.402917

**Authors:** Danny Salem, Yifeng Li, Pengcheng Xi, Hilary Phenix, Miroslava Cuperlovic-Culf, Mads Kaern

## Abstract

Accurate and efficient segmentation of live-cell images is critical in maximising data extraction and knowledge generation from high-throughput biology experiments. Despite recent development of deep learning tools for biomedical imaging applications, great demand for automated segmentation tools for high-resolution live-cell microscopy images remains in order to accelerate the analysis. YeastNet dramatically improves the performance of non-trainable classic algorithm, and performs considerably better than the current state-of-the-art yeast cell segmentation tools. We have designed and trained a U-Net convolutional network (named YeastNet) to conduct semantic segmentation on bright-field microscopy images and generate segmentation masks for cell labelling and tracking. YeastNet enables accurate automatic segmentation and tracking of yeast cells in biomedical applications. YeastNet is freely provided with model weights as a Python package on GitHub. https://github.com/kaernlab/YeastNet

## Background

*S. cerevisiae*, hereafter referred to as yeast, is a eukaryotic model organism used to study synthetic gene network development and analysis, as well as other biological processes. Bright-field and fluorescence microscopy are common approaches to investigate yeast behaviour. The quantitative analysis of time-lapse fluorescence microscopy images is a powerful tool for large-scale and single-cell analysis of dynamic and noisy cellular processes, such as gene expression (1–3). In an experimental design wherein yeast cells are expressing a fluorescent marker of interest, quantitative analysis involves the quantification of fluorescence intensity of pixels corresponding to cell regions in a microscopy image of those cells. Therefore, quantitative analysis requires accurate identification of cell regions, known as segmentation, as well as tracking of these cell regions between images, in the case of time-lapse microscopy.

In recent years, automated light microscopes have been paired with commercial or lab-constructed microfluidics devices to conduct time-lapse analysis of cell cultures in long-term perfusion conditions. Microfluidics-enabled time-lapse fluorescence microscopy allows the study of dynamic cellular processes in a single-cell manner. Automated image capturing of multiple fields of view at high imaging frequencies increases the number of tracked cells and enhances the resolution of the data. However, it also greatly increases the number of images that need to be analyzed from hundreds to tens of thousands. Thus, automated solutions for the problem of cell segmentation are necessary in order to avoid very time-consuming and error prone manual segmentation.

One way to address the segmentation problem has been the use of fluorescent markers expressed in the cytosol to label cells in fluorescence images. For example, cells with constitutively expressed green fluorescent protein (GFP) are easily separable from a background when imaged with a green light filter. Cell segmentation in fluorescence images is easier than that in corresponding bright-field images, however the number of different fluorescent proteins (FPs) that can be used in an experiment is limited (4). Due to overlapping excitation and emission spectra of many FPs, as well as the effects of Förster resonance energy transfer (FRET), using multiple different FPs requires careful selection. Moreover, additional fluorescent imaging increases the risk of phototoxicity and photobleaching. Although use of fluorescent markers can be beneficial in some cases, cell segmentation from bright-field images is required in the majority of cases. Therefore, the ideal algorithm for cell segmentation should work automatically from a bright-field image input.

Several features of yeast cells in bright-field images have been used for segmentation by non-trainable (manually parameterised) algorithms or manual analysis. A cell’s outline is generally its most distinguishable 2D structure in a bright-field image due to the distinct brightness of the outline relative to the cell interior and background. In slightly out of focus bright-field images, this outline is thicker and easier to identify using thresholding techniques (5). However, several characteristics of yeast cells prevent accurate segmentation with these traditional algorithms, particularly in the case of time-lapse microscopy wherein distinguishable features of cell outlines usually change over time. Yeast cells divide quickly, forming colonies of tightly packed cells within hours. In such dense colonies neighbouring cells converge or overlap, making detection of individual outlines very difficult. Additionally, cell outlines may change in brightness and thickness due to focus drift of the microscope over time.

### Related Work

Many algorithms have been presented for the cell segmentation problem over the past decade. Unsupervised algorithms which use computer vision processes like the watershed algorithm, image thresholding, and active contour fitting are popular solutions (6, 7). More recently, hybrid algorithms using pipelines of multiple processes with manual user supervision have become more common. One of these methods called CellStar uses thresholding and active contour fitting, in addition to user-enabled automated parameter fitting. CellStar shows the highest segmentation accuracy for a variety of data sets (8).

With the introduction of the convolutional neural net (CNN) design paradigm for the task of image classification (9–11), the deep learning approach for biomedical image analysis became very popular. Deep learning approaches for cell segmentation have been developed for various cell types and imaging methodologies (12–14), however, an accurate model is still lacking for general yeast cell segmentation in bright-field images. DeepCell, developed in 2016, is a deep learning model based on CNNs developed to segment bacterial and mammalian cells, but it has not been utilized for yeast cells. In 2017, a model trained to analyze yeast cells was published (12) based on the SegNet (15) architecture. Notably, this model was trained to detect yeast cells from very noisy differential interference contrast (DIC) microscopy images and can not be directly used for bright-field image analysis. In 2019, a Mask-RCNN model named YeastSpotter (16) was published for yeast cell segmentation. YeastSpotter is a model trained on the BBBC038 (17) dataset from the Broad Bioimage Benchmark Collection for a Kaggle competition for segmenting nuclei in mammalian cells. For a comparison with our approach, this model was taken and used directly on yeast bright field microscopy images without fine-tuning.

In recent years, the emergence of fully CNN for semantic segmentation has led to new computer vision possibilities in the field of biomedical imaging. Semantic segmentation is the process of segmenting an image into multiple classes by classifying each pixel in an image. The fully convolutional network for semantic segmentation (18) is the seminal architecture that introduced the ability to train a network to output pixel-wise class predictions based on every pixel in an input image. The fully connected layers usually found at the end of CNNs utilized to classify images were replaced with learned convolutional up-sampling layers. By using up-sampling layers, the down-sampling of the encoder layers is reversed and the output of the network maintains the resolution of the input image. In order to improve the up-sampling layer, feature maps obtained at earlier stages of the network are appended to the final layer via skip connections. It has been discovered that the increase in the number of connections leads to higher accuracy of the network (18).

In 2015, the U-Net architecture (19) improved classification accuracy achieved with the fully convolutional semantic segmentation networks introduced by Long et al. (18). The U-Net was developed for the task of pixel-wise cell segmentation of mammalian cells. By implementing skip connections for every layer in the down-sampling half of the network, each up-sampling layer consisted of twice the data. The U-Net demonstrated that it was not necessary to have very large annotated datasets to achieve high accuracy in segmentation tasks using deep learning.

The contribution of this work is two-fold. 1) We provide a dataset of 150 bright field images of budding yeast at three levels of focus, with ground truth segmentations. We also provide ground truth segmentation for 80 images from two datasets found in the Yeast Image Toolkit. This type of training data for bright field images of yest is very limited in this field, and will enable future research in this domain. 2) We present YeastNet, a deep learning model and tool that is shown to outperform the current state-of-the-art method for yeast cell segmentation. YeastNet is trained on images at multiple levels of focus to ensure invariance to shifting levels of focus in time-lapse experiments, a common problem with high-throughput microscopy experiments. It is the first time that a fully convolutional model has been applied in the domain of general bright field microscopy analysis of budding yeast.

## Results

We compared the segmentation performance of YeastNet on our dataset to a non-trainable (manually parameterized) classic algorithm adapted from a yeast cell-cycle research article (20). We also compared YeastNet to CellStar (8), the current state-of-the-art tool for bright-field yeast cell segmentation. We used the CellStar package and followed the instructions, to complete cell segmentation and tracking of our dataset. We also compared with YeastSpotter (16), a mask-RCNN deep learning model trained for the segmentation of mammalian cell nuclei. YeastNet was trained on just training portions of our dataset and was used to demonstrate the ability of this model to generalise to unseen test samples of our datasets and all samples in YIT Datasets 1 and 3. Furthermore, as a separate experiment, our model was also trained on a combination of training splits of all three datasets (we name this trained model YeastNet2) and was tested on test sets of the three datasets. The mean results obtained from 10-fold cross-validation are reported in Table 1. Clearly, our method achieves the highest cell IoU on our dataset. Our method also generalises as well or better as other methods, despite using a single dataset for training. Our model trained on training splits of all three datasets shows improved performance on all datasets. Fig. 1A shows a visual comparison of the segmentations by the non-trainable method, CellStar, and YeastNet. (As indicated in Table 1, YeastSpotter performed worse than CellStar, thus its segmentation is not provided here.)

**Table 1.** 10-fold cross validation of segmentation performance of our methods and benchmarks in terms of cell IoU.

| Method | Our Dataset | YIT Dataset 1 | YIT Dataset 3 |
| --- | --- | --- | --- |
| Non-trainable | 0.564 | 0.585 | 0.553 |
| CellStar | 0.699 | **0.693** | 0.693 |
| YeastSpotter | 0.561 | 0.609 | 0.677 |
| YeastNet | **0.878** | 0.687 | **0.726** |
| YeastNet2 | **0.886** | **0.820** | **0.853** |

**Fig. 1.**
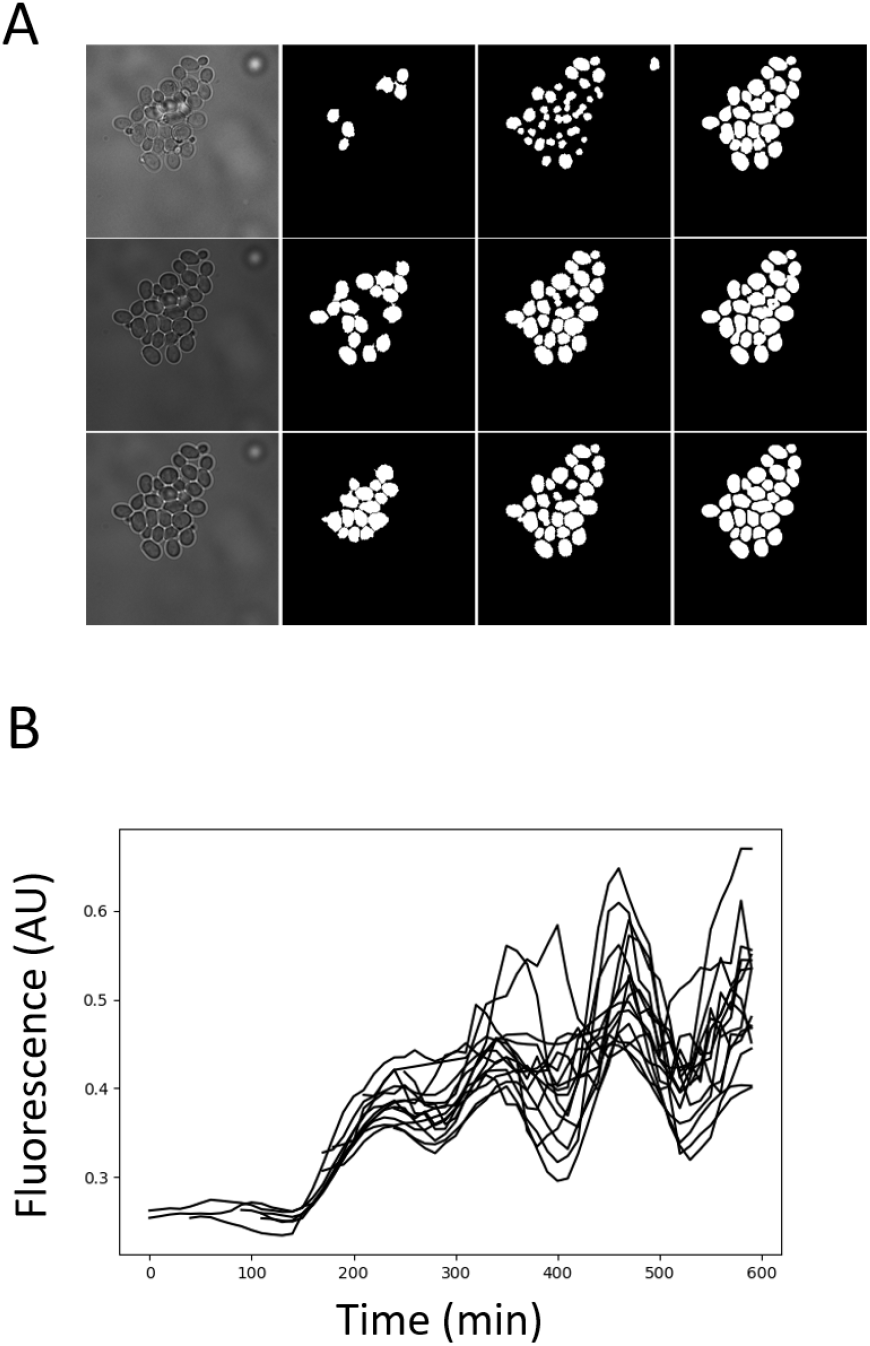
(A) Cell segmentation mask of the same colony at different levels of focus. From top to bottom the input images shown are: in focus, slightly out of focus, more out of focus. From left to right: cropped input image, and segmentations using the non-trainable method, CellStar and YeastNet. (B) Single cell fluorescence tracking, generated by YeastNet, to study dynamic gene expression. Each curve is a time-lapse mean pixel fluorescence measure describing the protein abundance of a dynamically expressed gene in an individually tracked cell.

**Fig. 2.**
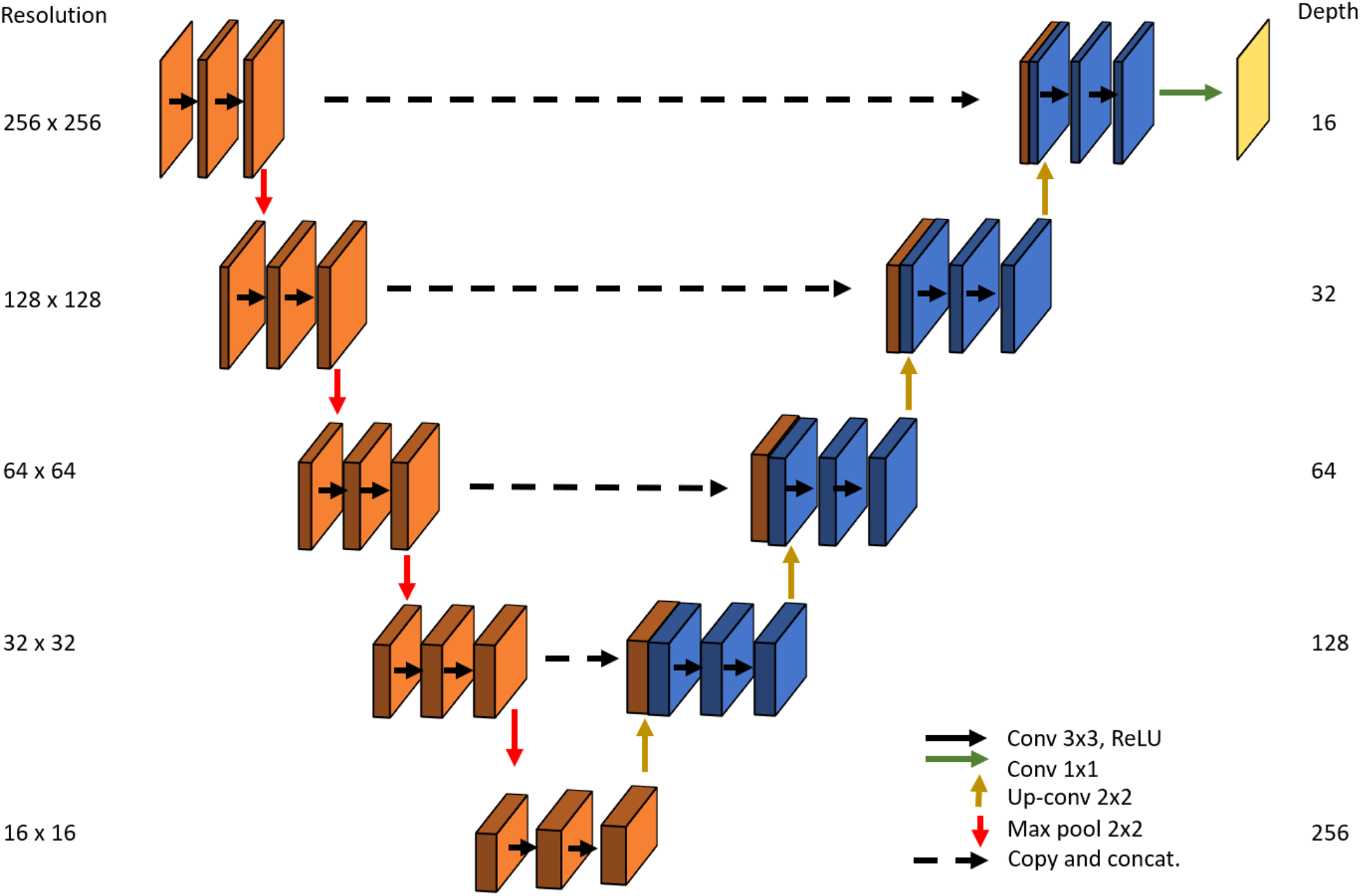
A visual description of the modified U-Net architecture applied in this work. Images in the down-sampling stage are orange and images in the up-sampling stage are blue. The pixel-wise class prediction output is yellow. Solid horizontal arrows indicate convolutional operations that do not change the height or width of the inference tensor. Vertical arrows indicate operations that reduce or increase the size of the inference tensor. Skip connections are represented by a dashed black arrow. The resolution of each tensor is shown on the left, and the feature depth at each level is shown on the right. The depth corresponds to the layer depth of each individual tensor in the row, tensors that result from skip connections have twice the depth. The output depth corresponds to the number of classes in the classification problem, in our case the dimensions of the output is 256*×*256*×*2.

Our dataset contains three different bright-field images at each time point, each corresponding to a different level of focus. We compared the segmentation and tracking performance of YeastNet and CellStar separately for each level of focus. We intended to train a model to be invariant to focus level by training on all types of images (images of the same colony at different levels of focus are treated as separate training examples). To test the tracking performance, it is necessary to use the entire set of images from a time-lapse, and invariably some of these images could come from the training set. To avoid this problem, a YeastNet3 model was trained using only transformed (rotated and mirrored) images. The test set consisted of only the original images with three levels of focus.

Table 2 shows both segmentation and tracking performances tested on the original time-course images at different levels of focus. As expected, CellStar’s segmentation performance is higher on out-of-focus images (Focuses 2 and 3) in comparison with its performance on in-focus images (Focus 1). YeastNet however has a higher cell IoU at both levels of focus, compared to CellStar. On the in-focus images, CellStar performs poorly, attaining less than a 0.5 cell IoU.

**Table 2.**
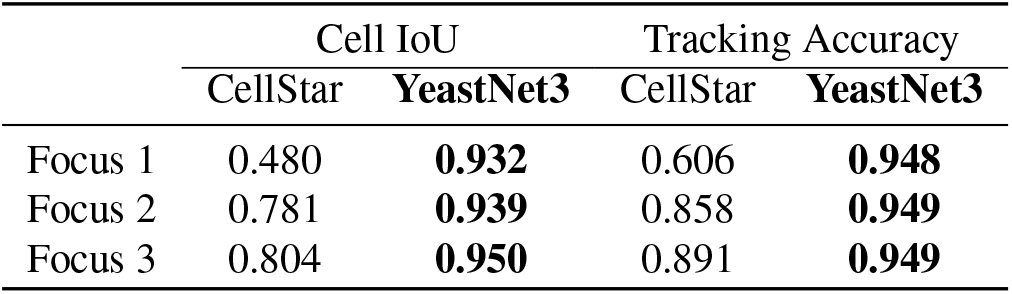
Cell tracking performance at different levels of focus. Focus 1: in focus, Focus 2: slightly out of focus, Focus 3: more out of focus.

|  | Cell IoU |  | Tracking Accuracy |  |
| --- | --- | --- | --- | --- |
|  | CellStar | **YeastNet3** | CellStar | **YeastNet3** |
| Focus 1 | 0.480 | **0.932** | 0.606 | **0.948** |
| Focus 2 | 0.781 | **0.939** | 0.858 | **0.949** |
| Focus 3 | 0.804 | **0.950** | 0.891 | **0.949** |

Meanwhile, YeastNet obtains a cell IoU that is almost double with 0.932. In addition, CellStar incorrectly segments air bubbles, as seen in Figure 1A, which YeastNet correctly ignores. YeastNet also maintains nearly the same segmentation accuracy with in-focus images as with out-of-focus images, while CellStar is over 15% lower. Thanks to its segmentation performance, YeastNet3 attains a higher tracking accuracy at every level of focus in comparison with CellStar, with the largest improvement (by 56%) at focus level 1. With the increased segmentation and tracking accuracy of YeastNet, the ultimate goal is to generate the plots as in Fig. 1B from cell tracks. Each curve describes single-cell time-lapse fluorescence over a 6 hour period.

## Discussion

In this work, we present a learnable U-Net model for the segmentation of yeast cells in bright-field images. Recent advances in this field have led to the development of very accurate tools for this task but they each have several trade-offs of varying severity. Among these trade-offs are: manual and time-consuming user input, unfeasible number of z-stack image for time-lapse analysis, and expensive equipment. We demonstrate that a deep learning approach to this problem can yield an accurate yeast cell segmentation tool that requires minimal user input, minimal data per time point, and can be used with common and widely available imaging platforms. The high performance attained by YeastNet3 at all three levels of focus indicates that it is resistant to the segmentation errors usually caused by changes in focus. Resistance to changes in focus is a very important feature because focus drift is a significant problem that can cause many problems with image analysis (21), especially with large experiments where hundreds or thousands of images are being taken every hour.

There are several challenges in applying computer vision to this domain. A common problem with time-lapse microscopy is drift in the focus of images. The focus of bright-field images is very important because nearly all segmentation algorithms rely on the yeast cells being at a certain focal length for accurate segmentation. By using the 3 different z-stacks of each time point as separate training examples, our YeastNet model learned to detect yeast cells at different focal lengths; in essence making the trained model invariant to minor changes in the focus. A qualitative comparison of the performance on different focal lengths between our model and CellStar is shown in Fig. 1A and Table 2. The higher segmentation performance leads to an increased tracking accuracy. Furthermore, the wider difference in the IoU between CellStar and YeastNet is significant for downstream applications of the cell traces. Due to variation in fluorescent protein within the cell, accurate fluorescence quantification requires entire cells to be segmented.

We also present a new, labeled dataset for training computer vision models to segment bright-field microscopy images. Since annotated data of this kind was not publicly available, only data we labelled manually could be used to train this model. Manually segmenting images to generate the datasets is laborious and due to the limited size of the dataset, we discovered that minor biases in what constitutes a cell can have many downstream effects. There is some ambiguity in what is and isn’t considered part of a cell, especially at different levels of focus. A computer vision model will learn to detect cells in the same way that cells are segmented in the ground truth. Therefore, comparisons conducted with ground truth segmentations created using different guidelines for what constitutes part of a cell, will lead to poor reported performance, even if performance appears qualitatively very high. To train a robust model, more labelled data is required. This will enable the model to generalize better to new types of datasets, including data from: different imaging modalities, lighting conditions, resolution, and magnification levels. Analysis of time-lapse fluorescence m icroscopy u sually involves cell segmentation followed by cell tracking. In this work, we used a simple linear sum assignment solution to the problem of cell tracking since our focus was to develop and train a network for yeast cell segmentation. With this simple solution, YeastNet still achieves very high tracking performance. An adaptation of a recent algorithm for cell tracking like the algorithm used in (8), or a new deep learning approach would lead to even higher tracking performance. Furthermore, there has been recent developments in U-Net design, with variations like U-Net++ (22), and the 3DU-Net (23). The improvement of YeastNet through the use of improved U-Net designs will also be explored in the next version of YeastNet.

## Conclusion

We designed YeastNet to improve the accuracy of identifying individual *S. cerevisiae* cells from bright-field microscopy images. Our model is based on the U-Net semantic segmentation architecture and it was trained using a manually labelled dataset. YeastNet segments bright-field images by generating pixel-wise predictions between background and cell classes. We compared YeastNet to a classic method and the current state-of-the-art tool for general *S. cerevisiae* segmentation and tracking. We achieve higher performance in every metric used: intersection over union, segmentation accuracy, and tracking accuracy.

We also present a new dataset, consisting of 150 bright-field microscopy images of budding yeast. This dataset consists of 50 fields of view of a growing colony, taken at 3 levels of focus, as well as, manually segmented ground-truth segmentation masks. This is the first publicly available dataset for bright-field microscopy segmentation of yeast, and we hope it is used to advance research in this domain.

## Methods

### Datasets

Bright-field microscopy images of yeast were produced in house as part of analysis of novel reporter model. Yeast cells were designed to include synthetic gene networks that allow user regulated production of the reporter. Fluorescence time-lapse microscopy was done to study the yeast cells under different regulatory regimes using an inverted light microscope with an automated stage and focus control. The images taken at each time point and colony are: 3 bright-field images at different focus level (for segmentation) and 2 fluorescence images (for expression level quantification). Single colonies from antibiotic plates were inoculated in synthetic media (1% adenine, 2% glucose). Overnight cultures were diluted to logarithmic phase (OD_600_ of 0.1). Cell cultures were diluted to an OD_600_ of 0.07 prior to loading of the chamber. Diluted cultures were loaded into CellASIC ONIX Y03C microfluidic plates for microscopy analysis. The ONIX software loading protocol (8 psi for 15 seconds) was used for loading. ten fields of view were located containing a single colony of 1-3 cells. The cells were trapped due to the height of the microfluidics chamber and grew in a monolayer. Colonies did not reach confluence and rogue cells did not pass through the field of view.

Imaging was controlled by Nikon NIS-Elements software. Colonies were imaged every 10 minutes. Each field of view was imaged several times per imaging cycle: a GFP fluorescence image, an mCherry fluorescence image, and bright field images at 3 levels of focus (in focus, 0.6*μ*m above focal plane, 1.2*μ*m above focal plane). Three bright field images at and around the centre of cells were taken to enable cell segmentation and to counteract focal drift in the Nikon PFS auto-focus system. Images were captured with a CoolSnap HQ2 camera connected to a Nikon Ti-E inverted microscope with a 60x oil immersion objective. Images were taken at a resolution of 1340 × 1092 with an exposure time of 200ms. The experiment lasted 10 hours, or 60 timepoints. Only the first fifty time points were manually segmented to generate a ground-truth dataset. Each of the 50 time points include 3 bright-field images captured of the same colony. Thus, the 50 ground truths create a dataset of 150 available input images with corresponding labelled true segmentations. Manual segmentation was performed in MATLAB using the CTseed algorithm provided in Doncic et al. (7), which allows the user to manually select a segmentation threshold and manually correct segmentation errors. In addition to increasing the sample size, using the 3 image captures (i.e. z-stacks) as separate training images enables the model to be less reliant on the focus of the image, a notorious problem of cell segmentation algorithms (21).

In addition to our dataset, images were also taken from two datasets in the Yeast Image Toolkit. The Yeast Image Toolkit (YIT) was created by the authors of Cellstar(8); they created the ground truth labels for the images in the YIT datasets which were provided by the Batt & Hersen lab (24). Images were taken every 3 minutes with a 50ms exposure. A 100x oil immersion objective was used with Olympus IX81 inverted microscope. Images were taken at a resolution of 512 × 512. Cells were fixed a nd g rew i n a m onolayer and did not reach confluence. Y IT D ataset 1 c onsisted o f 60 frames with a cell count starting with 14 cells and growing to 26 cells. YIT Dataset 3 consisted of 20 frames with a cell count starting with 101 cells and growing to 128 cells. These datasets were chosen due to their qualitative and quantitative differences. The associated ground truth for these datasets does not include cell masks so we generated ground truth masks using ImageJ (25). Additional details can be found on the Yeast Image Toolkit website. (http://yeast-image-toolkit.org/pmwiki.php)

To standardize images from different datasets prior to network inference, images were normalized on a dataset-level. The mean and standard deviation of the training set for an individual dataset were taken and used to zero center and normalize the images in the test and training set images of that dataset. Images were also re-scaled between 0 and 1.

### Data Augmentation

It is important to employ several techniques to augment the limited dataset which was manually labelled. Doing so improves the training of the model and enhances its generalisation capacity to unseen data. Yeast cells are generally ellipse-shaped, and their orientation is not important. Rotating and flipping the training images increase the size of the dataset and could improve the model’s invariance to orientation. This also increases the ability to accurately classify debris in the background as background. Debris looks different in different experiments and increasing the variety of debris used in images is important for generalisation to other datasets of this domain. Furthermore, random cropping was used to increase the variety in the training images, but also to decrease the memory required for storing a training image. Using 256 × 256 crops of the 1024 × 1024 microscopy images enabled the use of larger batch sizes during training.

### Non-Trained Method

The yeast cell segmentation algorithm in (20) uses two out of focus bright-field images, one below and one above the focal plane, to generate a cell segmentation mask. It then uses a CFP fluorescence microscopy image to correct for false positives. We adapted and modified t his a lgorithm t o w ork w ith a s ingle o ut o f f ocus image, above the focal plane, and removed the false positive correcting functionality since it relied on an extra imaging modality. Parameters were manually optimized for each experiment. Parameters included minimum and maximum cell size, watershed and thresholding parameters, and optimal cell circularity.

### Proposed Segmentation Model

We designed a convolutional network based on the U-Net semantic segmentation architecture (19) which is a fully convolutional network that combines an encoder network that generates a dense feature map, followed by a decoder network that generates pixel-wise classification p redictions. F ig. 2 is a visual description of the model developed in this work. By zero-padding the tensor for every convolution operation, the output prediction tensor maintains the same size dimensions as the input image.

The network is composed of repeated motifs that consist of two 3×3 convolutional + ReLU layers and a resolution changing layer. In the encoder part of the network, the resolution changing layer in each motif is a 2×2 max-pooling layer. Four repeats of this down-sampling motif make up the encoder network. The decoder follows with four repeats of an up-sampling motif whose final layer is a transpose convolution. The final set of operations consists of two 3 3 convolutional + ReLU layers and a 1×1 convolutional layer which predicts the class probability for each pixel.

### Weighted Loss Function

In cell segmentation, there is a very important spatial class imbalance issue that must be accounted for in any loss calculation. The number of background pixels separating cells is very small but their correct classification is crucial for accurate segmentation and cell labelling. Without proper weighting of pixel-wise loss, a classifier can attain a very low loss by simply learning to separate colonies from the background. A weighted loss function that emphasizes the accurate prediction of the areas between cells is crucial to train the semantic segmentation of cells from background.

Cross entropy is a very common loss function in machine learning and it has been adapted to computer vision problems as a pixel-wise calculation. To account for the spatial class imbalance, we use a weighted pixel-wise cross-entropy loss function, first described in (19). First, we generate a class imbalance weight matrix (*w*_*c*_) and scale it so that the weight for every cell is 1. Next, a weight matrix favouring pixels near multiple cells with higher weights is calculated using the following equation:

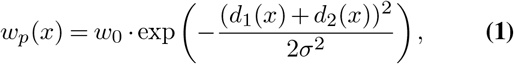

where *x* is the location of a pixel point; *d*(*x*) is defined as the distance from pixel *x* to the nearest pixel belonging to a cell, *d*_1_(*x*) and *d*_2_(*x*) are thus the distances in pixels from the current pixel to the nearest two cells; *w*_0_ and *σ* are manually set parameters (in our training process, they were both set to 10). To create the final weight map, the two weight matrices are added together,

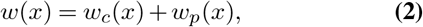

to get the final weight map for a training image.

The weight map for one of the training images is shown in Fig. 3. Each pixel is weighted by its proximity to cells. Therefore, the pixels between cells have a very high weight.

**Fig. 3.**
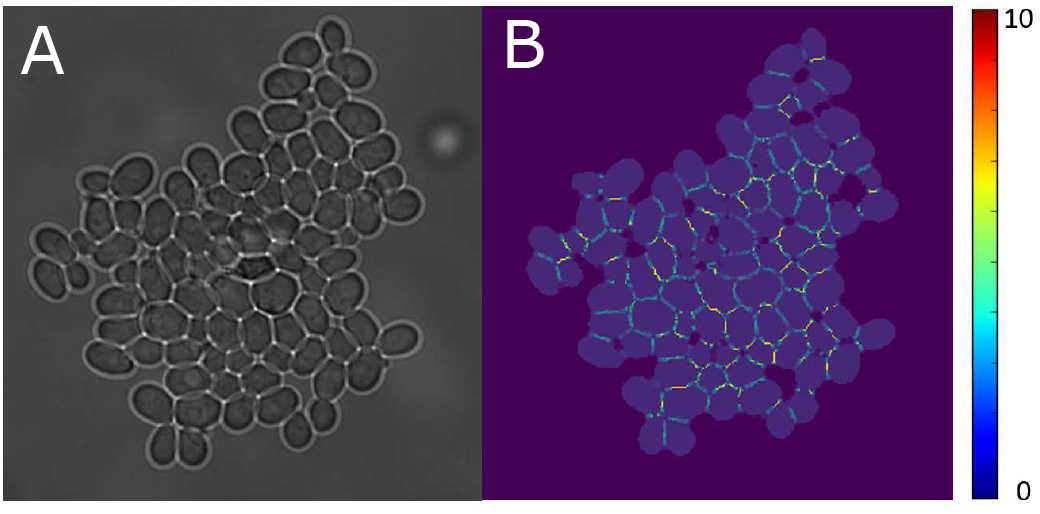
(B) The weighted loss matrix used to bias the pixel-wise cross-entropy loss and the (A) microscopy image it was generated for. The pixel color in (B) indicates the final weight taking into account class imbalance and proximity to cells.

### Cell Tracking

As cell tracking was not the main focus of this work, the algorithm used to create cell lineages is simple and effective. The problem of cell tracking is framed as a linear sum assignment optimization by calculating the combinatorial distances between all cell centroids in subsequent images. The optimization is solved using the Hungarian algorithm (26–28) which minimizes the sum of differences between all the paired centroids.

This algorithm is formally described for two frames (*t*, *t* + 1) using the equation:

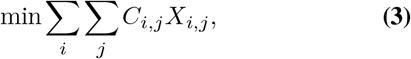

where *C* is a difference matrix describing the cost of pairing cell *i* in frame *t* and cell *j* in frame *t* + 1. In our case the cost is distance between the centroids of the two cells: *i* and *j*. *X* is a Boolean matrix holding the assignments between cells. *X*_*i,j*_ = 1 if *i* and *j* are assigned to be the same cell.

Tracking accuracy is defined using an F-measure statistic. The set of all true cell pairs between subsequent images is compared against the predicted cell pairs. The equation used for this metric is:

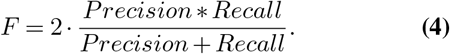

## Author’s contributions

Study Concept: DS, MK. Study Design and Approach: DS, YL, PX. Developed and tested Model: DS. Data Preparation: DS, HP. Manuscript Writing: DS, YL, PX, HP, MK, MC. All authors read and approved the manuscript.

## Acknowledgements

Not applicable

## Availability of data and materials

The datasets and code used during the current study are available at https://github.com/kaernlab/YeastNet. Two publicly available datasets were used from the Yeast Image Toolkit (Dataset 1 and Dataset 3). The Yeast Image Toolkit is available at: http://yeast-image-toolkit.biosim.eu/pmwiki.php

